# Diverse myeloid cells are recruited to the developing and inflamed mammary gland

**DOI:** 10.1101/2020.09.21.306365

**Authors:** Gillian J Wilson, Ayumi Fukuoka, Francesca Vidler, Gerard J Graham

## Abstract

The immune system plays fundamental roles in the mammary gland, shaping developmental processes and controlling inflammation during infection and cancer. Here we reveal unanticipated heterogeneity in the myeloid cell compartment during development of virgin, pregnant and involuting mouse mammary glands, and in milk. We investigate the functional consequences of individual and compound chemokine receptor deficiency on cell recruitment. Diverse myeloid cell recruitment was also shown in models of sterile inflammation and bacterial infection. Strikingly, we have shown that inflammation and infection can alter the abundance of terminal end buds, a key developmental structure, within the pubertal mammary gland. This previously unknown effect of inflammatory burden during puberty could have important implications for understanding the control of pubertal timing.

## Introduction

The mammary gland is a highly regenerative tissue within the body. It is unique, in that most of its development occurs postnatally throughout the female reproductive lifetime. The gland undergoes dramatic structural changes throughout puberty where proliferative structures, terminal end buds (TEB), invade through the surrounding fatty stroma, giving rise to a complex epithelial network in adulthood (MM Richert, KL Schwertfeger, JW Ryder, 2000). During pregnancy, rapid proliferation of epithelial cells generates lobuloalveoli (LAL), and milk producing ducts form during lactation. When lactation stops upon weaning, a process of involution occurs where 90% of the gland remodels to its pre-pregnancy form (MM Richert, KL Schwertfeger, JW Ryder, 2000).

The immune system has long been identified as a key component of the mammary gland, residing within a stromal population containing fibroblasts, extracellular matrix (ECM), and adipocytes (Wiseman and Werb, 2002). In particular, macrophages play an essential role in regulating mammary gland branching morphogenesis, as development is dramatically impaired in macrophage-deficient mice (Pollard and Hennighausen, 1994; Gouon-Evans, Rothenberg and Pollard, 2000). In addition, the density of branching is reduced in CCL11 deficient mice, which have decreased numbers of eosinophils (Gouon-Evans, Rothenberg and Pollard, 2000). Mast cell degranulation is also necessary for normal ductal development (Lilla and Werb, 2010). The adaptive immune system plays an inhibitory role in the regulation of pubertal development through CD11c+ antigen presenting cells and CD4+ T cells (Plaks *et al*., 2015). Neutrophils and dendritic cells are also key cell types present in the gland during involution (Atabai, Sheppard and Werb, 2007; Betts *et al*., 2018).

During inflammation, the immune cell landscape of the mammary gland is altered. In breast cancer, immune cells infiltrate the gland, including tumour associated macrophages (TAMs), Myeloid derived suppressor cells (MDSC), tumour associated neutrophils (TANs), T-cells, and NK cells (Nagarajan and McArdle, 2018). Mastitis is a common disease of the breast, caused by a build-up of milk in the ducts and exacerbated by bacterial infection. Increased numbers of leukocytes including neutrophils, monocytes and macrophages, are detected in the gland and in milk (Hassiotou *et al*., 2013; Cacho and Lawrence, 2017). This impacts milk quality, leading to reduced infant weight gain and dysregulated immune development (Tuaillon *et al*., 2017) and often leads to early cessation of breastfeeding.

The molecular mechanisms which regulate the movement of immune cells as they migrate within the gland to mediate their effects, are not fully understood. Insights into these processes will enhance our understanding of how immune cells contribute to mammary gland development and protect against inflammation. Chemokines, characterised by a conserved cysteine motif, are a family of proteins important in cell recruitment, and as in vivo regulators of intra-tissue cell movement. The chemokine family is comprised of CC, CXC, XC and CX3C sub-families according to cysteine distribution, and chemokines act through G-protein coupled receptors to facilitate leukocyte migration (Nibbs and Graham, 2013). Inflammatory chemokine receptors (iCCRs: CCRs1, 2, 3 and 5)) are often expressed by immune cells and are required for cell recruitment within the body (Douglas P. Dyer *et al*., 2019). Previously we have shown important roles for chemokine receptors in shaping the macrophage dynamics within the mammary gland to control pubertal development (Wilson *et al*., 2017, 2020).

Here we reveal unanticipated heterogeneity in myeloid cells within the mammary gland at key developmental stages in virgin, pregnant and involuting mice. We also reveal the myeloid cell composition of murine milk. In addition, we show that diverse myeloid cells are recruited to the mammary gland during local inflammation and remote infection. Importantly, we have shown that inflammation and infection alter the number of terminal end buds, a key developmental structure, within the pubertal mammary gland. The direct effect of inflammatory burden on pubertal development has not been reported previously and could have important implications for understanding the control of pubertal timing.

## Results

### Leukocyte levels in the mammary gland throughout development

Flow cytometry was carried out to determine the levels of immune cells in the mammary gland at key time points throughout virgin development, pregnancy and involution, and in maternal milk. Leukocytes were defined as CD45+ and gated as outlined in Supplementary Figure 1. The percentage of CD45+ cells within the live population significantly increased between early (5 weeks) and late (6.5 weeks) puberty (Figure 1ai). CD45+ cells represent a much lower percentage of live cells within the gland on the first day of involution and in milk. The absolute number of CD45+ cells per gland (Figure 1aii) was found to peak during late puberty at 7 weeks in the virgin mammary gland and at day 15.5 during pregnancy. A detailed flow cytometric analysis was carried out to identify myeloid cell types using a defined panel of cell surface markers (Table 1, Supplementary Figure 1, 2). t-SNE analysis carried out of CD45+ cells revealed distinct clusters corresponding to the cell types identified (Figure 1 b).

**Table 1:** Myeloid cells within the mammary gland.

|  | Surface Marker |  |  |  |  |  |  |  |
| --- | --- | --- | --- | --- | --- | --- | --- | --- |
| Cell Type | CD11b | CD11c | F4/80 | SiglecF | Ly6C | Ly6G | MHCII | CD206 |
| CD11b+F4/80+<br>Macrophage | + |  | + |  |  |  |  |  |
| CD206+<br>macrophage |  |  | + | - |  |  |  | + |
| CD11b+CD11c+<br>macrophage | + | + | + |  |  |  | + |  |
| Ductal<br>macrophage | - | + | + |  |  |  | + |  |
| CD11b-CD11c-<br>macrophage | - | - | + |  |  |  | + |  |
| CD11b+CD11c-<br>macrophage | + | - | + |  |  |  | + |  |
| Ly6C low<br>monocyte | + |  |  |  | low | - |  |  |
| Ly6C high<br>monocyte | + |  |  |  | high |  |  |  |
| Eosinophil-1 | + |  | + | + |  |  |  |  |
| Eosinophil-2 | + |  | - | + |  |  |  |  |
| Neutrophil | + |  | - |  |  | + |  |  |
| Dendritic cell |  | + | - |  |  |  | + |  |
| G-MDSC | + |  | + |  |  | + |  |  |

**Figure 1:**
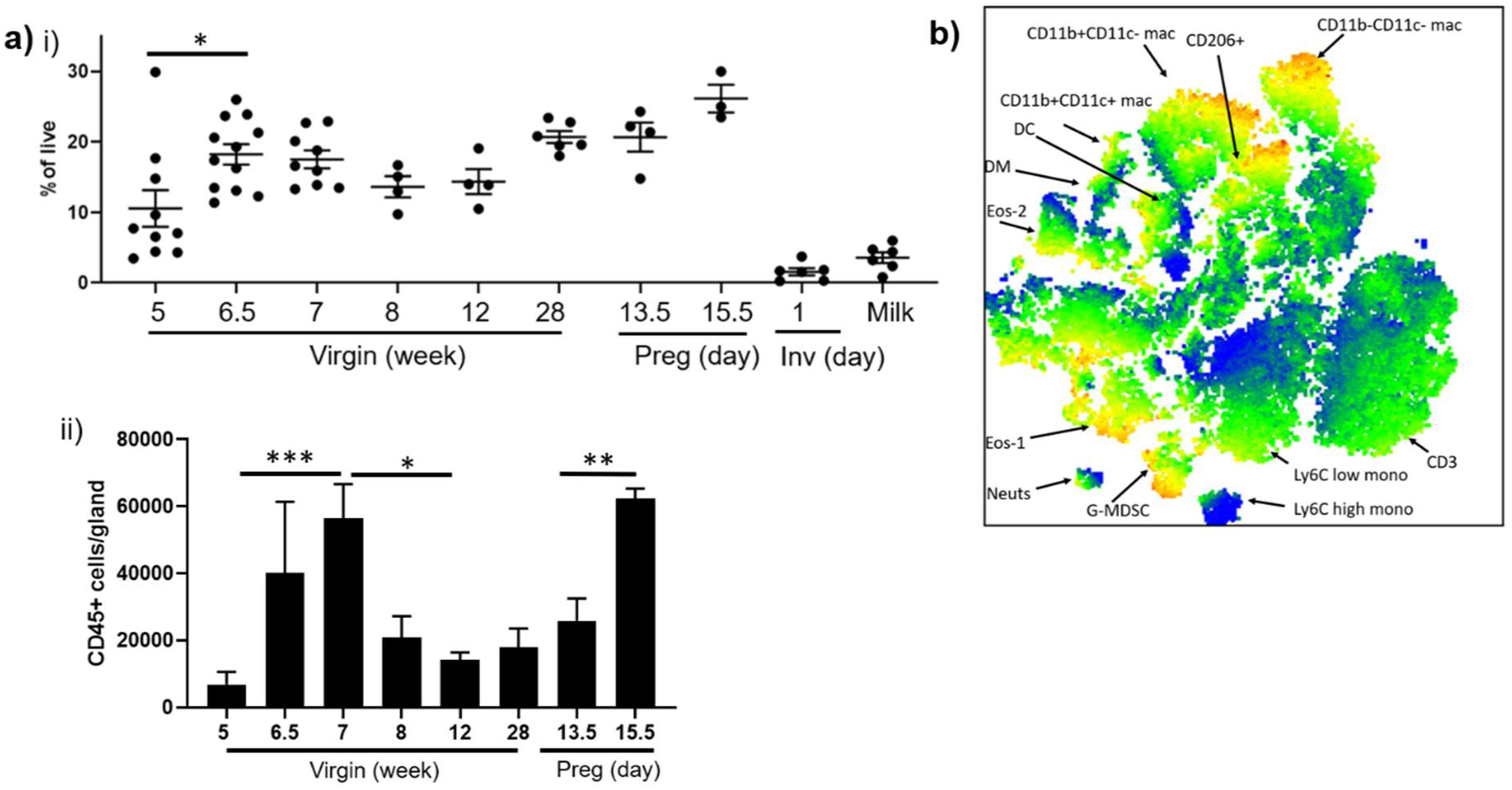
Leukocyte levels in the mammary gland throughout development. **a)** Flow cytometry of CD45+ cells in the mammary gland expressed as **i)** a percentage of live cells during virgin development (5 weeks, n=10, 6.5 weeks, n=12, 7 weeks, n=9, 8 and 12 weeks, n=4, 28 weeks n=6) pregnancy (day 13.5, n=4, day 15.5, n=3) involution (n=6) and in milk (n=6). **ii)** Total CD45+ cells per gland, during virgin development (5 weeks, n=10, 6.5 weeks, n=3, 7 weeks, n=6, 8 and 12 weeks, n=4, 28 weeks n=6), and pregnancy (day 13.5, n=4, day 15.5, n=3). **b)** Representative tSNE analysis of CD45+ cells within a sample from day 13.5 of pregnancy. Significantly different results are indicated. Error bars represent S.E.M.

### Diverse macrophage subsets in the mammary gland throughout development

Macrophages play key roles in promoting mammary gland development and protecting against disease. Classically mammary gland macrophages have been defined by CD11b and F4/80 positivity. However recent studies have revealed further complexity in the macrophage populations within the mammary gland. We identified a key population of SiglecF-F4/80+CD206+ macrophages recruited by CCR1 which promote branching morphogenesis during puberty (Wilson *et al*., 2020). In addition, Dawson *et al* identified a novel ductal macrophage important in tissue remodelling, defined as CD11b-CD11c+MHCII+F4/80+(Dawson *et al*., 2020). Here we reveal that in addition to these subsets, there are three further populations, CD11b+CD11c+, CD11b+CD11c-and CD11b-CD11c-macrophages (Table 1). The gating strategy employed is shown in Supplementary Figure 1. We investigated the presence of each of these subtypes in the gland throughout development and in milk.

CD11b+F4/80+ or ‘classic’ macrophages increase during puberty between 5 and 6.5 weeks (Figure 2 a). They represent a substantial proportion (approx. 20-30%) of the CD45+ cells within the mammary gland through adulthood (8 weeks to 6 months) and pregnancy. During early involution of the mammary gland and in breast milk they represent only around 5% of CD45+ cells. This population can be further subdivided into CD11c-MHCII+ (CD11b+CD11c-), and CD11c+MHCII+ (CD11b+CD11c+) macrophages. CD11b+CD11c-macrophages are present at each stage of virgin development and pregnancy, at a reduced level in early involution, but not in milk (Figure 2 b). CD11b+CD11c+ macrophages are a small population which significantly increase during puberty, between 5 and 6.5 weeks, and again during aging, between 12 and 28 weeks (Figure 2 c). They are detected at low levels during pregnancy, involution and in milk (Figure 2 c).

**Figure 2:**
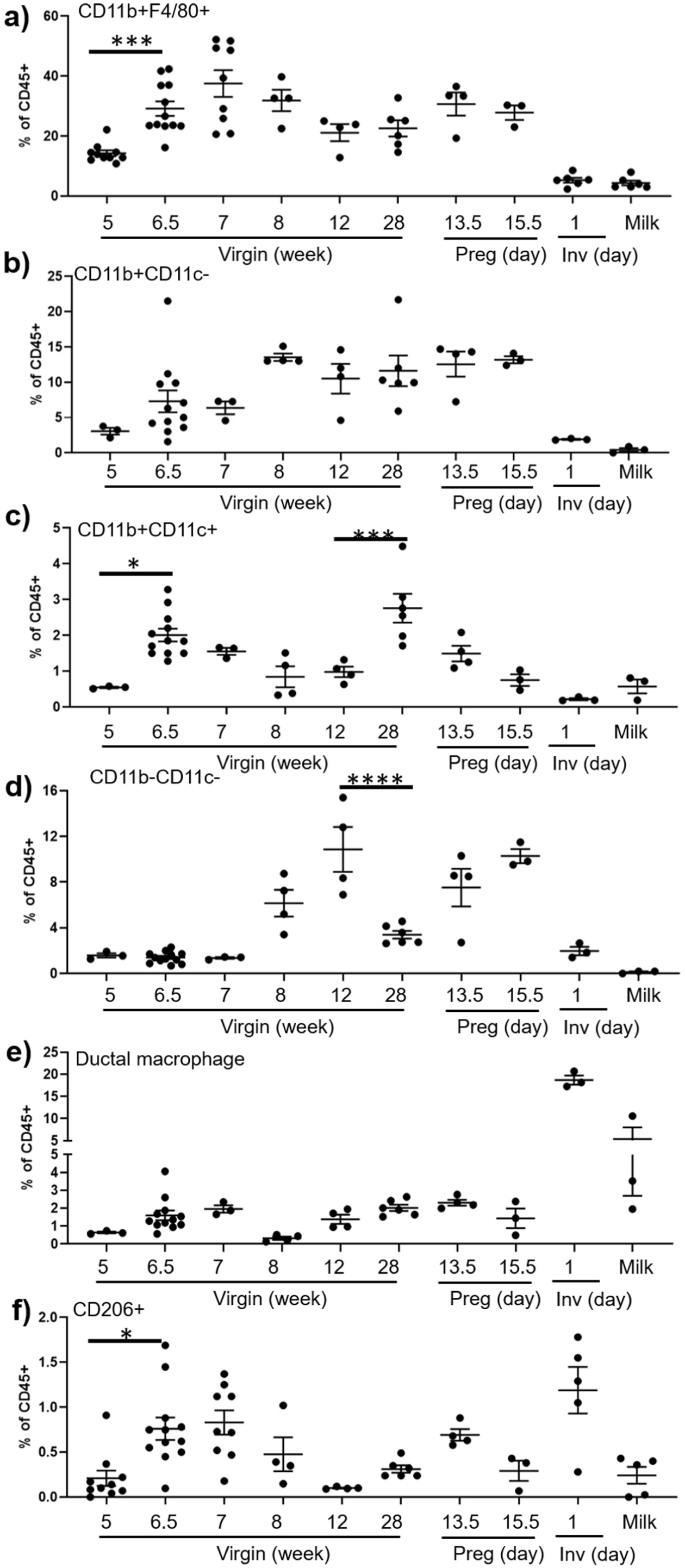
Diverse macrophage subsets in the mammary gland throughout development. Flow cytometry was used to determine the percentage of **a)** CD11b+F4/80+ macrophages, **b)** CD11b+CD11c-MHCII+F4/80+ macrophages, **c)** CD11b+CD11c+MHCII+F4/80+ macrophages, **d)** CD11b-CD11c-MHCII+F4/80+macrophages and **e)** CD11b-CD11c-MHCII+F4/80+ ductal macrophages, **f)** SiglecF-F4/80+CD206+ macrophages within the CD45+ compartment of the mammary gland, during virgin development, pregnancy, involution and in milk. **a, f)** virgin (5 weeks, n=10, 6.5 weeks, n=12, 7 weeks, n=9, 8 and 12 weeks, n=4, 28 weeks n=6) pregnancy (day 13.5, n=4, day 15.5, n=3) involution (n=6) and milk (n=6). **b-e)** virgin (5 weeks, n=3, 6.5 weeks, n=12, 7 weeks, n=3, 8 and 12 weeks, n=4, 28 weeks n=6) pregnancy (day 13.5, n=4, day 15.5, n=3) involution (n=3) and milk (n=3). Significantly different results are indicated. Error bars represent S.E.M.

We also identified a population of CD11b-CD11c-macrophages throughout development which are found at the highest levels in early adulthood and pregnancy, but not in milk (Figure 2 d). Ductal macrophages (CD11b-CD11c+) represent a low percentage of CD45+ cells throughout virgin development and pregnancy. However, in early involution they comprise around 20% of CD45+ cells (Figure 2 e). CD206+ macrophages represent a small proportion of CD45+ cells but have an important role promoting branching morphogenesis in puberty (Wilson *et al*., 2020). Here we show a significant increase in this population between early (5 weeks) and late puberty (6.5 weeks) (Figure 2 f). They are also detected in the gland throughout adulthood, pregnancy and involution, and in milk (Figure 2 f).

### Monocytes, granulocytes and dendritic cells in the mammary gland

We next examined the presence of further myeloid cell types in the mammary gland using the gating strategy outlined in Supplementary Figure 2 and Table 1. There are 2 populations of monocytes in the mammary gland, Ly6C high and Ly6C low, which increase in late puberty (Figure 3 a, b). Both monocyte subtypes are also found in adult virgin and pregnant glands, at low levels during early involution and at higher levels in milk (Figure 3 a, b). There are also 2 distinct populations of eosinophils characterised by F4/80 expression. Type 1 eosinophils (F4/80+) represent a higher percentage of CD45+ cells than type 2 (F4/80-) (Figure 3c, d). The percentage of type 1 but not type 2 eosinophils increases during late puberty. Both are detected in adult, pregnant and involuting glands, and in milk (Figure 3c, d). Neutrophils are present at low levels throughout virgin and pregnant gland development, but increase during involution and represent a substantial proportion, around 15%, of leukocytes in milk (Figure 3 e). Similarly, we have identified small numbers of dendritic cells (DCs) throughout virgin and pregnant development, which dramatically rise during involution (Figure 3 f). DCs are also detected in milk (Figure 3 f). A small population of granulocytic myeloid derived suppressor cells (G-MDSC) was also detected at each of the key developmental stages in the mammary gland, and in milk (Figure 3 g).

**Figure 3:**
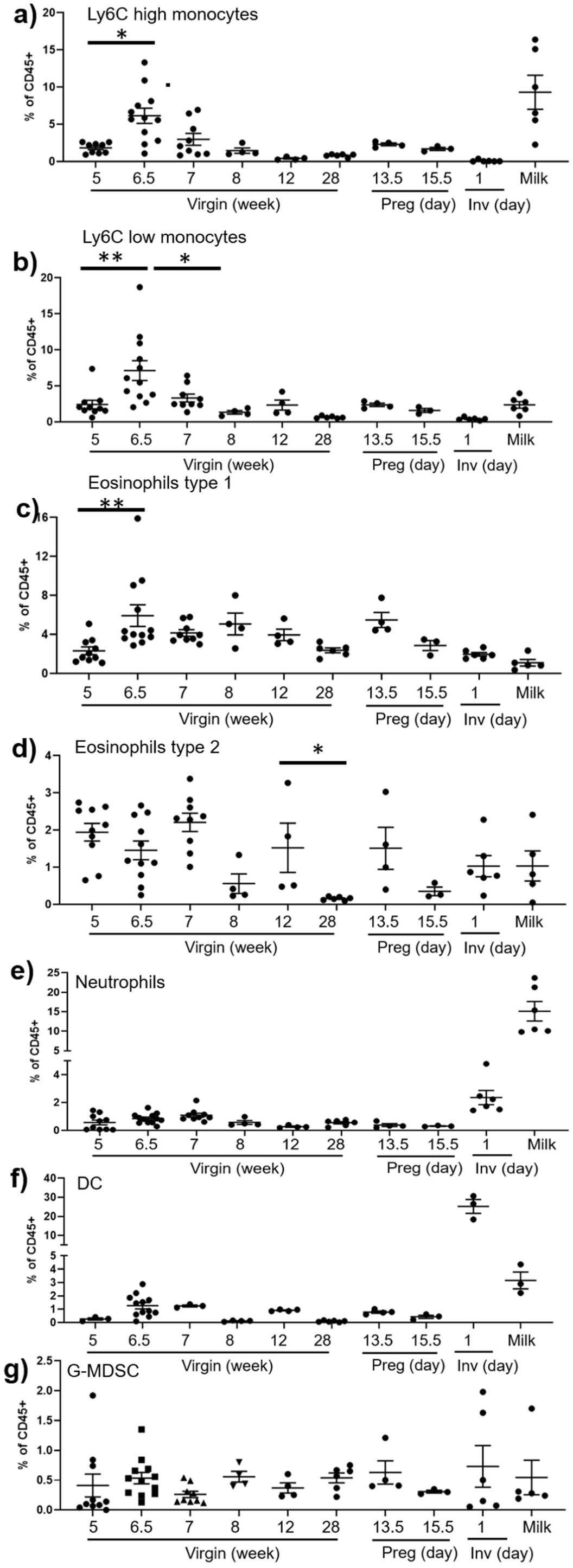
Monocytes, granulocytes and dendritic cells in the mammary gland throughout development. Flow cytometry was used to determine the percentage of **a)** CD11b+Ly6C high monocytes **b)** CD11b+Ly6C low Ly6G-monocytes, **c)** Type 1 CD11b+SiglecF+ F4/80+ eosinophils **d)** Type 2 CD11b+SiglecF+ F4/80-eosinophils **e)** F4/80-CD11b+Ly6G+ neutrophils, **f)** F4/80-CD11c+MHCII+ dendritic cells, and **g)** F4/80+CD11b+Ly6G+ granulocytic myeloid derived suppressor cells (G-MDSC) within the CD45+ compartment of the mammary gland, during virgin development, pregnancy, involution and in milk.. **a-e, g)** virgin (5 weeks, n=10, 6.5 weeks, n=12, 7 weeks, n=9, 8 and 12 weeks, n=4, 28 weeks n=6) pregnancy (day 13.5, n=4, day 15.5, n=3) involution (n=6) and milk (n=6). **f)** virgin (5 weeks, n=3, 6.5 weeks, n=12, 7 weeks, n=3, 8 and 12 weeks, n=4, 28 weeks n=6) pregnancy (day 13.5, n=4, day 15.5, n=3) involution (n=3) and milk (n=3). Significantly different results are indicated. Error bars represent S.E.M.

CSF1R is a marker for cells of the mononuclear phagocyte lineage including macrophages, monocytes, and granulocytes (Sasamono et al, 2003). We carried out flow cytometry of *MacGreen* (CSF1R GFP reporter) transgenic mice to confirm that each of the cell types we have identified in the mammary gland are of myeloid origin (Supplemental Figure 3 a). Surprisingly, we also found a small percentage of CD45 negative cells that are CSF1R positive (Supplemental Figure 3 b). Confocal imaging and flow cytometric analysis reveal CSF1R expression by epithelial cells (Supplemental Figure 3 b).

### iCCRs are required for cell recruitment to the mammary gland

To investigate which iCCRs are required for the recruitment of myeloid cells to the mammary gland, we analysed key cellular populations in WT, individual receptor deficient mice, and mice with a compound receptor deletion of all four iCCRs, CCRs1, 2, 3 and 5. The receptor deficient mouse strains are on different genetic backgrounds and therefore have been compared with their appropriate WT. Previously we revealed that CD206+ macrophages are reduced in CCR1 deficient mice during late puberty (7 weeks)(Wilson *et al*., 2020). There are no significant differences in any of the other myeloid cell populations investigated in this study (Figure 4 a). In the absence of the key monocyte receptor CCR2, both Ly6C high and Ly6C low monocytes are depleted in 7 weeks old mice (Figure 4 b). We also observed a reduction in type 1 eosinophils (Figure 4 b). Previously we observed that CD11b+F4/80+ cells were unaffected in adult CCR2-/-mice (Wilson *et al*., 2017), however in this study we observe a reduction in CD11b+F4/80+ cells during puberty (Figure 4 b). In CCR3-/-mammary glands there is a significant reduction in Type 2 eosinophils but not in any of the other populations investigated. (Figure 4 c). In the absence of CCR5 we observed no differences in cell recruitment (Figure 4 d). Importantly, in pubertal iCCR-/-mice which lack all 4 receptors, we also observed significant reductions in Ly6C high and Ly6C low monocytes, type 1 and 2 eosinophils, and CD11b+F4/80+ macrophages. This recapitulates the results of the individual receptor deficient mice and suggests there are no additional combinatorial effects of compound receptor deficiency.

**Figure 4:**
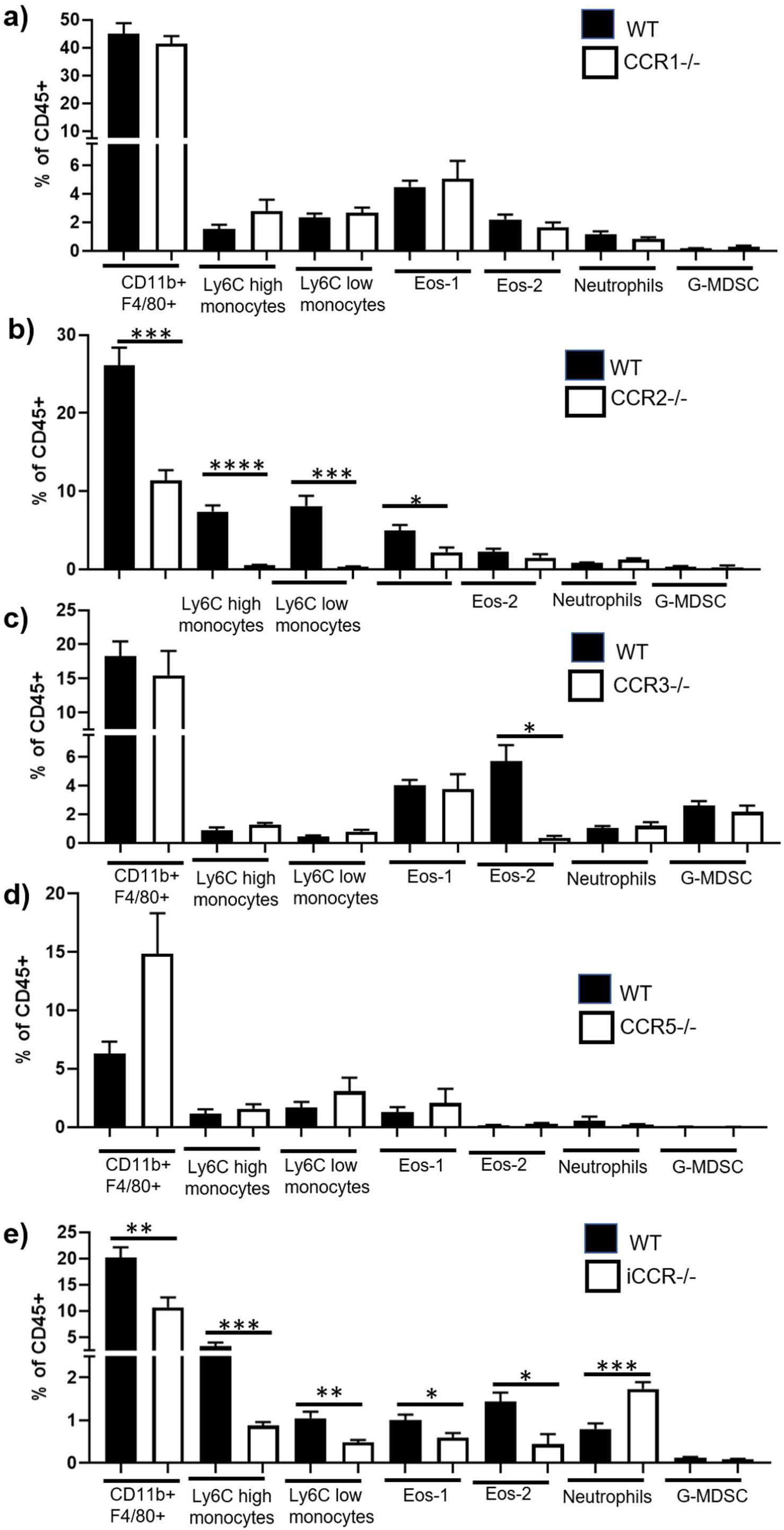
iCCRs are required for cell recruitment to the mammary gland. Flow cytometry was used to determine the percentage of CD11b+F4/80+ macrophages, Ly6C high monocytes, Ly6C low monocytes, type 1 (eos-1) and 2 (eos-2) eosinophils, neutrophils and G-MDSC within the CD45+ compartment of the mammary gland during virgin development of **a)** WT and CCR1-/-(7 weeks, n=6 per group), **b)** WT (7 weeks, n=11) and CCR2-/-(7 weeks, n=6), **c)** WT and CCR3-/-(7 weeks, n=4 per group), **d)** WT and CCR5-/-(12 weeks, n=4 per group), **c)** WT and iCCR-/-(7 weeks, n=7 per group). Significantly different results are indicated. Error bars represent S.E.M.

### Myeloid cells are recruited to the mammary gland during inflammation and infection

To investigate the immune response in the mammary gland during local inflammation, we injected mice during late puberty (6-7 weeks), subcutaneously at the site of the mammary gland with either PBS or 500 µg of FITC labelled *Escherichia coli* particles (ECP) for 18 h and 5 days. Cell recruitment to the mammary gland was measured by flow cytometry. Overall, the number of CD45+ cells significantly increased 18 h after challenge with ECP, with no difference observed in the resolution phase, after 5d (Figure 5 a). Specifically, neutrophils, Ly6C high and Ly6C low monocytes, CD11b+F4/80+ macrophages, type 1 eosinophils and G-MDSCs are all increased in the mammary gland 18 h after challenge with ECP (Figure 5 a). In addition, we observed binding of the FITC labelled ECP to each of these cell types (Figure 5 b). After 5 days the number of cells was not significantly different and bound FITC labelled ECP were not detected (Figure 5). We observed no change in recruitment of, or ECP binding, to ductal CD11b+CD11c+, CD11b+CD11c-, CD11b-CD11c-or CD206+ macrophages, type 2 eosinophils or dendritic cells.

**Figure 5:**
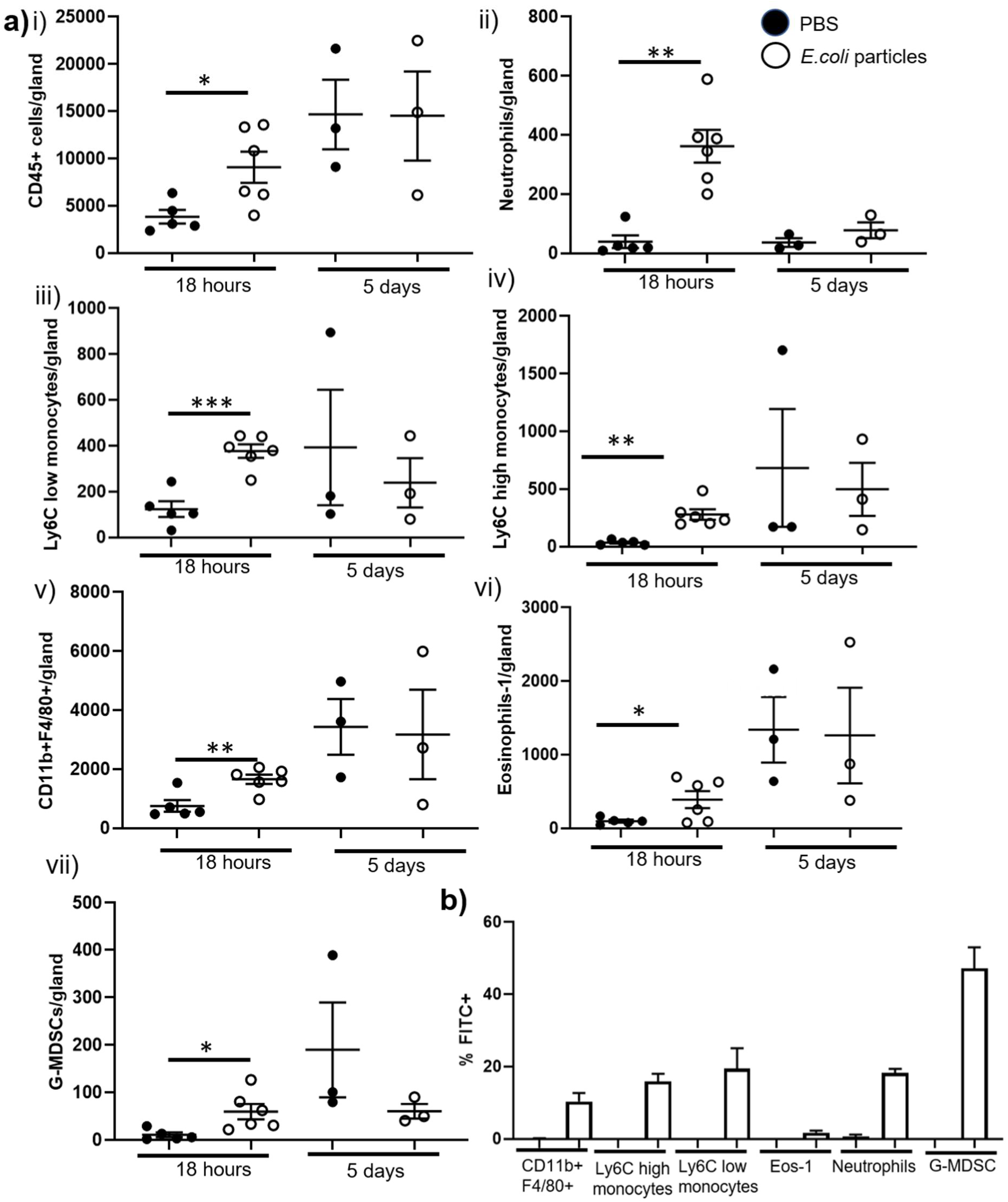
Myeloid cells are recruited to the mammary gland during sterile inflammation. Flow cytometry was used to determine **a)** the number of **i)** CD45+ cells, **ii)** neutrophils, **iii)** CD11b+F4/80+ macrophages, **iv)** Ly6C low monocytes, **v)** Ly6C high monocytes, **vi)** type 1 eosinophils (eos-1), and **vii)** G-MDSC within the mammary gland after challenge with either PBS (denoted by black circles) or 500 µg of FITC labelled *E*.*coli* particles (white circles), for 18h (PBS, n=5, *E*.*coli* particles, n=6) and 5 days (n=3 per group). **b)** The percentage of cells bound by FITC labelled *E*.*coli* particles 18h after challenge (PBS, n=5, *E*.*coli* particles, n=6). Significantly different results are indicated. Error bars represent S.E.M.

To investigate whether myeloid cells are altered in the mammary gland during a live bacterial infection, 6-7 week old mice were infected intraperitoneally with 1 × 10^6^ CFU of a uropathogenic strain of *E*.*coli* (CFT073). 5 days after infection of the peritoneum, the number of CD45+ cells within the mammary gland increased (Figure 6 a). As observed during sterile inflammation, increased numbers of neutrophils, Ly6C low monocytes and CD11b+F4/80+ macrophages were observed (Figure 6 b-d). We also detected increased numbers of dendritic cells and CD206+ macrophages during infection, which was not seen during sterile challenge (Figure 6 e-f).

**Figure 6:**
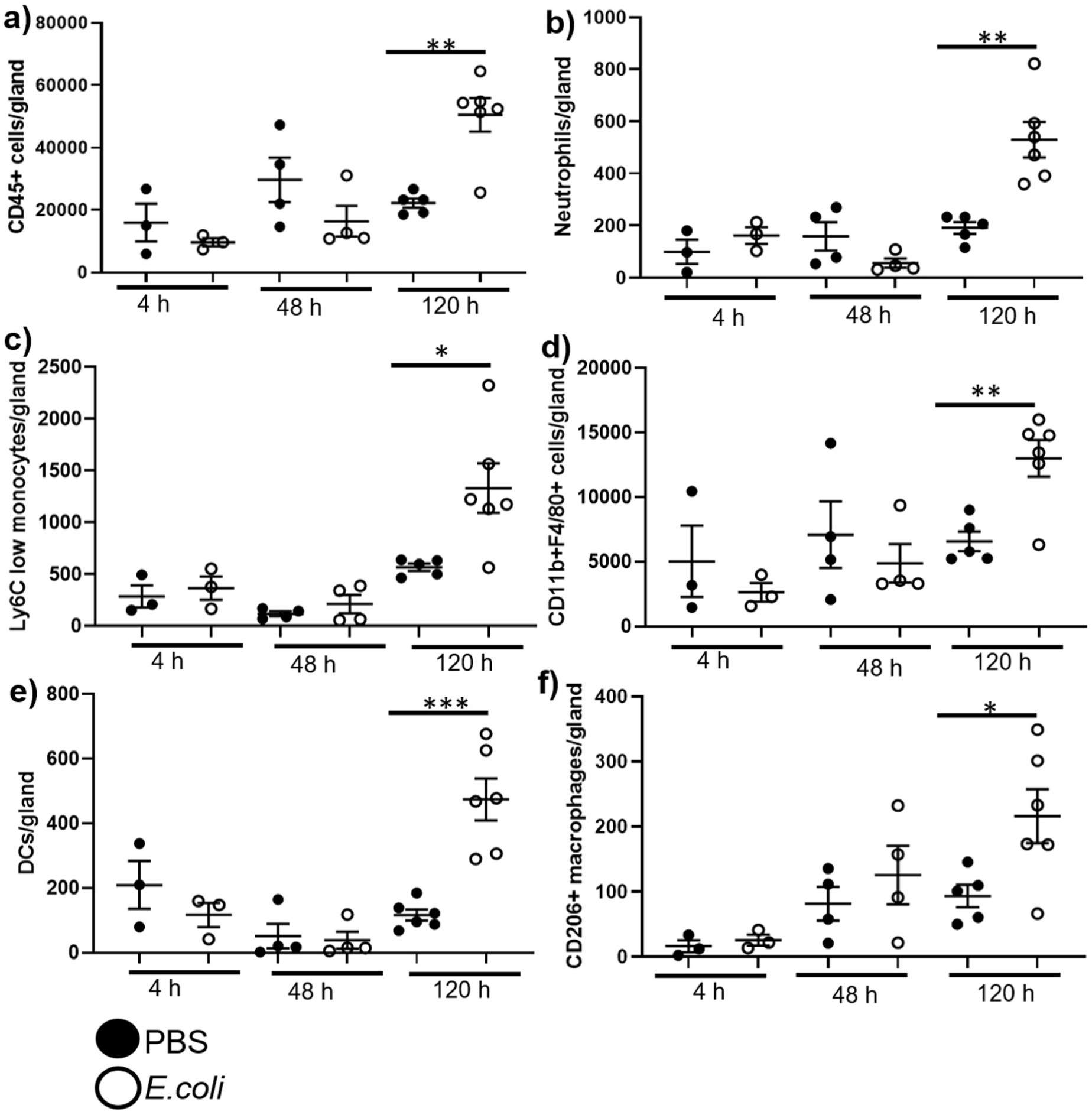
Myeloid cells are recruited to the mammary gland during infection. Flow cytometry was used to determine the number of **a)** CD45+ cells, **b)** neutrophils, **c)** Ly6C low monocytes, **d)** CD11b+F4/80+ macrophages, **e)** Dendritic cells, and **f)** CD206+ macrophages within the mammary gland after intraperitoneal challenge with either PBS (denoted by black circles) or 1x 10^6^ CFU *E*.*coli* strain CFT073 (white circles), for 4 h (n=3 per group), 2 days (n=4 per group), and 5 days (PBS, n=5, *E*.*coli*, n=6). Significantly different results are indicated. Error bars represent S.E.M.

### Mammary gland structures are altered during infection and inflammation

We next investigated whether inflammation of the mammary gland altered development of the gland during puberty. We analysed carmine alum stained whole-mounts of mammary glands from pubertal WT mice challenged with either PBS, 200 µg ECP or 1 × 10^6^ CFU *E*.*coli* CFT073 (Figure 7). TEBs are highly proliferative structures within the pubertal mammary gland which give rise to the ductal epithelial network within the gland. Strikingly we found that 3 d after intravenous challenge with 200 µg of ECP, the average number and width of terminal end buds was markedly reduced (Figure 7 a). We also investigated the effect of live bacterial infection of the peritoneum on TEB formation in the mammary gland and found that TEBs were reduced 2 and 5 days after infection (Figure 7 bi). However, the morphology of the TEB was not affected during peritoneal infection (Figure 7 bii).

**Figure 7:**
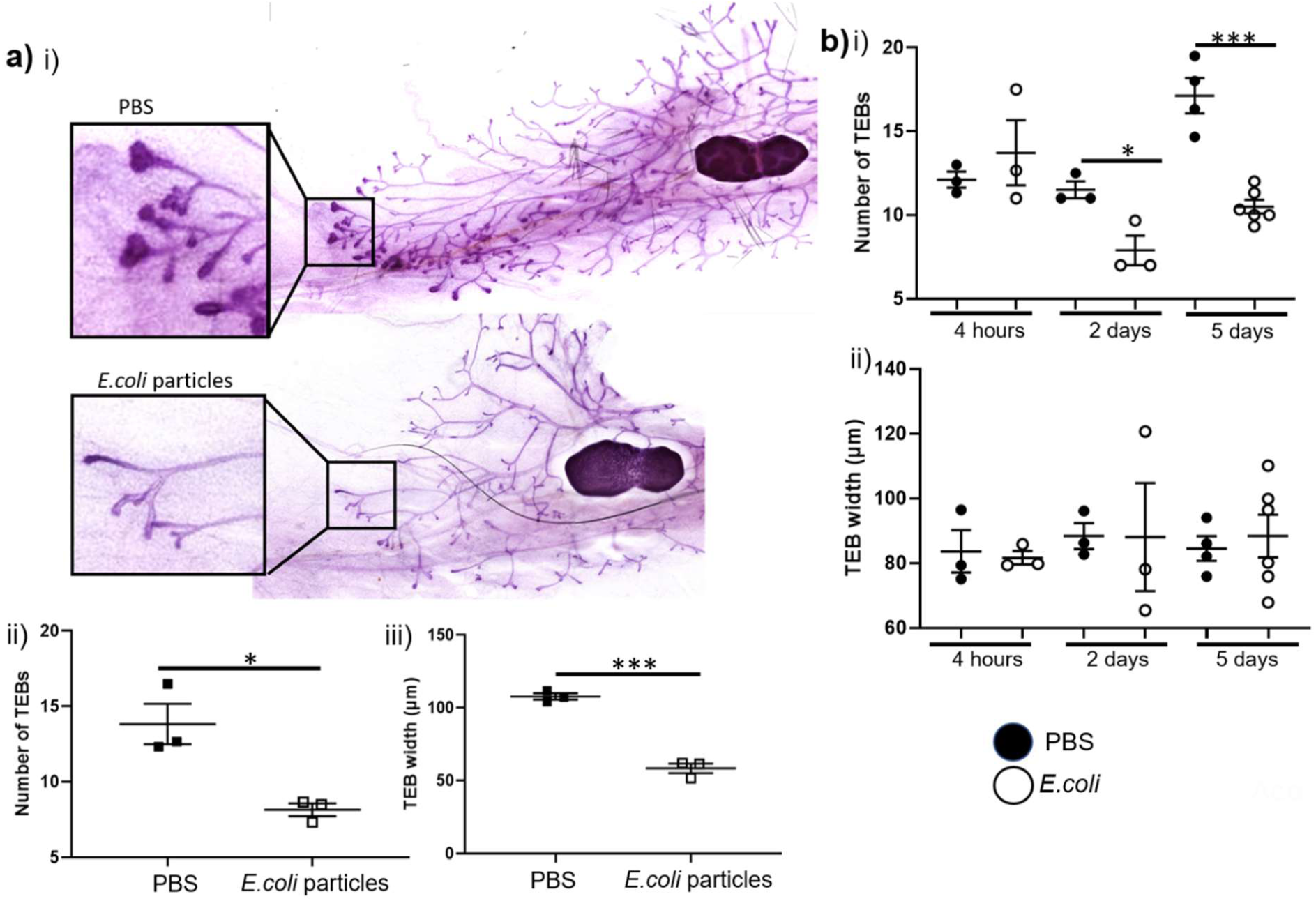
Mammary gland structures are altered during infection and inflammation. **a) i)** Representative carmine alum whole mount images (10 x) of pubertal virgin mammary glands 3 days after intravenous challenge with PBS or 200 µg *E*.*coli* particles (ECP). **ii)** The number of terminal end buds (TEB) was determined as the average number from at least 2 individual fields of view (FOV) (5×) per gland and **iii)** the average width of all TEBs was determined from at least 2 F.O.V (5x) per gland, 3 d after i.v challenge with PBS (denoted by black squares) or ECP (white squares) (n=3 per group). **b) i)** The number of TEBs and **ii)** average width of all TEBs after intraperitoneal challenge with either PBS (black circles) or 1x 10^6^ CFU *E*.*coli* strain CFT073 (white circles), for 4 h (n=3 per group), 2 days (n=3 per group), and 5 days (PBS, n=4, *E*.*coli*, n=6). Significantly different results are indicated. Error bars represent S.E.M.

## Discussion

Previously our understanding of immune system diversity within the mammary gland in normal development has been limited. Here we provide a detailed analysis of the myeloid cell landscape in the mouse mammary gland at key time points in development and in murine milk. Macrophages are a particularly heterogeneous cell type within the mammary gland. Recent studies have revealed novel subsets such as the CD206+ macrophages which promote branching in late puberty (Jäppinen *et al*., 2019; Wilson *et al*., 2020), and ductal macrophages which cover the epithelium in lactation and involution to facilitate remodelling (Dawson *et al*., 2020). Here we have used multi-parameter flow cytometry to reveal a further three macrophage subsets within the mammary gland CD11b+CD11c+, CD11b+CD11c- and CD11b-CD11c-. Further studies to investigate whether these macrophages have overlapping, or distinct functions could reveal important insight into macrophage control of mammary gland development and surveillance.

In addition, we have identified other important myeloid cell types in the mammary gland. We believe this study represents the first full investigation of myeloid cells throughout each of the key developmental stages. Although not exclusively, a high proportion of mammary gland macrophages, at rest, and in inflammation, are derived from monocytes in the bone marrow (Coussens and Pollard, 2011). Here we reveal that both Ly6C high and Ly6C low monocytes are present in the developing mammary gland at each stage, increasing significantly between early and late puberty. In the absence of CCR2 both subtypes are depleted in the mammary gland. Monocytes are also identified in established murine milk and likely contribute to neonatal immune protection. We have also defined 2 types of eosinophil based on F4/80 positivity. Type 1 (F4/80+) eosinophils only increase in late puberty and are recruited through a CCR2 dependent mechanism. In contrast, type 2 (F4/80+) eosinophils are recruited through CCR3 and are unchanged in puberty. To our knowledge, our study is the first to describe the presence of small populations of neutrophils, G-MDSC and DCs in the mammary gland of mice during virgin and pregnant development. As has been reported previously, neutrophils are found at higher numbers in early involution and are found at high levels in milk (Atabai, Sheppard and Werb, 2007; Cacho and Lawrence, 2017).

We also carried out a detailed analysis of myeloid cell recruitment to the gland in individual and compound iCCR deficient mice which lack all 4 receptors. We are able to confirm that CCR2 is the dominant monocyte receptor in the mammary gland, as has been observed throughout the body (Douglas P. Dyer *et al*., 2019). Notably, reduced recruitment of myeloid cells in iCCR-/-mice corresponded with the cell types reduced in individual receptor deficient mice. This suggests that there are no additional combinatorial effects of multiple receptor deficiency.

Importantly, in this study we have revealed that local subcutaneous inflammation and bacterial infection at a distant site with the body can have profound effects on the immune cells present in the mammary gland. For the first time we have been able to show that an inflammatory environment directly affects developmental structures within the mammary gland. TEBs drive the formation of the ductal epithelial network during puberty. Here we have shown that during inflammation and infection the number of TEBs decreases. It has been shown that early breast development leads to higher risks of breast cancer in later life (Bodicoat *et al*., 2014), and women with dense epithelial networks in the breast are more likely to develop breast cancer (Nazari and Mukherjee, 2018). Thus, the potential to manipulate the immune system to delay branching in puberty could have significant health benefits. There is evidence to suggest that increased infections in early life lead to a delay in female puberty (Kwok *et al*., 2011). This previously unknown effect of inflammatory burden on mammary structures during puberty could have important implications for understanding how pubertal timing is controlled.

## Methods

### Animals

Animal experiments were carried out under a UK Home Office Project Licence and conformed to the animal care and welfare protocols approved by the University of Glasgow. C57BL/6 mice, ACKR2-/-(Jamieson *et al*., 2005), CCR1-/-, CCR3-/-, CCR5-/-and iCCR-/-(Douglas P Dyer *et al*., 2019), and *MacGreen* mice (Sasmono *et al*., 2003) were bred at the specific pathogen-free facility of the Beatson Institute for Cancer Research. All mice used for experiments in this study were female.

### Mammary gland digestion

The inguinal lymph node was removed, and the fourth inguinal mammary gland was chopped coarsely. Enzymatic digestion with 3 mg/ml collagenase type 1 (Sigma) and 1.5 mg/ml trypsin (Sigma) was carried out in a 37°C shaking incubator at 200 rpm for 1 h, in 2 ml Leibovitz L-15 medium (Sigma). Tissue was shaken for 10 s before 5 ml of L-15 medium supplemented with 10% foetal calf serum (Invitrogen) was added. Centrifugation was carried out at 400 g for 5 min. Red Blood Cell Lysing Buffer Hybri-Max (Sigma) was applied for 1 min prior to washing in PBS containing 5 mM EDTA. Cells were then resuspended in 2 ml 0.25% Trypsin-EDTA (Sigma) and incubated for 2 min at 37°C. Next 5 ml of L-15 containing 1 μg/ml DNase1 (Sigma) was added for 5 min at 37°C. L-15 containing 10% FCS was then added to stop the reaction and cells were filtered through a 40 μm cell strainer. Finally, cells were washed in FACS buffer (PBS containing 1% FCS and 5 mM EDTA). Milk was obtained by removing mammary glands from lactating mice at day 7. Mammary glands were placed intact in FACS buffer and milk was collected, washed and stained.

### Flow cytometry

Antibodies were obtained from BioLegend and used at a dilution of 1:200: CD45 (30-F11), CD11b (M1/70), F4/80 (BM8), SiglecF (S17007L), Ly6C (HK1.4), CD11c (N418), MHCII (M5/114.15.2), EpCAM (G8.8), CD49f(GoH3), and CD206 (C068C2) for 30 min at 4°C. Ly6G (1A8) was obtained from BD Bioscience. Dead cells were excluded using Fixable Viability Dye eFluor 506 (Thermo Fisher). Flow cytometry was performed using a Fortessa, (BDBiosciences) and analysed using FlowJo V10.

### Sterile Inflammation of the mammary gland

Female mice between 6-7 weeks of age were injected subcutaneously with 500 µg, or intravenously with 200 µg of FITC labelled *E*.*coli* (K-12 strain) Bioparticles in 200 µl PBS (Thermo). After a defined number of days, mice were culled and mammary glands were excised and processed for whole mount and cellular analysis.

### Peritoneal bacterial infection

Female mice between 6-7 weeks of age were injected intraperitoneally with 1×10^6^ CFU of *E*.*coli* (CFT073 strain). Bacteria were grown overnight in Luria-Bertani medium, before being sub-cultured and grown to log phase for injection (OD_600_ = 0.5, 5×10^8^ CFU/ml). Mice were monitored for weight loss and clinical signs of infection. Mice were culled and mammary glands were excised and processed for whole mount and cellular analysis.

### Carmine Alum Whole Mount

Carmine alum whole mounts were carried out as previously described (Wilson *et al*., 2017, 2020). Briefly, fourth inguinal mammary glands were fixed in 10% neutral buffered formalin (NBF) (Leica) overnight at 4°C. Glands were dehydrated for 1 h in distilled water, then 1 h 70% ethanol and 1 h 100% ethanol before incubation in xylene overnight (VWR international). Rehydrated was achieved by 1 h incubation in 100% ethanol, 70% ethanol and distilled water, before staining with Carmine Alum solution at room temperature overnight (0.2% (w/v) carmine and 10 mM aluminium potassium sulphate (Sigma)). Tissue was dehydrated again and incubated overnight in xylene. Glands were then mounted with DPX (Leica) and 10× magnification stitched bright-field images were obtained using an EVOS FL auto2 microscope (Thermofisher). 5 x brightfield images were obtained using the Zeiss Axioimager M2 with Zen 2012 software. TEBs were counted as the average from at least 2 F.O.V. from each whole mount. All samples were blinded before measurements were taken.

### Statistical analysis

Data were analysed using GraphPad Prism 8.1.2. Normality was assessed using Shapiro Wilk and Kolmogorov–Smirnov tests. For normally distributed data, two-tailed, unpaired t-tests were used. Where data was not normally distributed, Mann–Whitney tests were used. Multiple comparison analysis was carried out using an ANOVA with Tukey’s post-test. Significance was defined as p<0.05 *. Error bars indicate standard error of the mean (S.E.M.).

## Author Contributions

GJW conceived the study, performed experiments, analysed data and wrote the paper. AF and FV performed experiments. GJG conceived the study and wrote the paper.

## Acknowlegments

We thank the University of Glasgow’s animal facility staff for the care of our animals and flow cytometry facility staff for technical assistance. We thank Prof. Andy Roe for providing *E*.*coli* CFT073. The study was supported by a Programme Grant from the Medical Research Council (MR/M019764/1). Work in GJG’s laboratory is also funded by a Wellcome Trust Investigator Award (<u>099251/Z/12/Z</u>). GJG is a recipient of a Wolfson Royal Society Merit award.

## References

Atabai, K., Sheppard, D. and Werb, Z. (2007) ‘Roles of the Innate Immune System in Mammary Gland Remodeling During Involution’, Journal of Mammary Gland Biology and Neoplasia, 12(1), pp. 37–45. doi: 10.1007/s10911-007-9036-6.

Betts, C. B., Pennock, N. D., Caruso, B. P., Ruffell, B., Borges, V. F. and Schedin, P. (2018) ‘Mucosal Immunity in the Female Murine Mammary Gland’, The Journal of Immunology, 201(2), pp. 734 LP – 746. doi: 10.4049/jimmunol.1800023.

Bodicoat, D. H., Schoemaker, M. J., Jones, M. E., McFadden, E., Griffin, J., Ashworth, A. and Swerdlow, A. J. (2014) ‘Timing of pubertal stages and breast cancer risk: the Breakthrough Generations Study’, Breast Cancer Research, 16(1), p. R18. doi: 10.1186/bcr3613.

Cacho, N. T. and Lawrence, R. M. (2017) ‘Innate Immunity and Breast Milk’, Frontiers in Immunology, 8, p. 584. doi: 10.3389/fimmu.2017.00584.

Coussens, L. M. and Pollard, J. W. (2011) ‘Leukocytes in mammary development and cancer’, Cold Spring Harbor perspectives in biology. Cold Spring Harbor Laboratory Press, 3(3), p. a003285. doi: 10.1101/cshperspect.a003285.

Dawson, C. A., Pal, B., Vaillant, F., Gandolfo, L. C., Liu, Z., Bleriot, C., Ginhoux, F., Smyth, G. K., Lindeman, G. J., Mueller, S. N., Rios, A. C. and Visvader, J. E. (2020) ‘Tissue-resident ductal macrophages survey the mammary epithelium and facilitate tissue remodelling’, Nature Cell Biology, 22(5), pp. 546–558. doi: 10.1038/s41556-020-0505-0.

Dyer, Douglas P., Medina-Ruiz, L., Bartolini, R., Schuette, F., Hughes, C. E., Pallas, K., Vidler, F., Macleod, M. K. L., Kelly, C. J., Lee, K. M., Hansell, C. A. H. and Graham, G. J. (2019) ‘Chemokine Receptor Redundancy and Specificity Are Context Dependent’, Immunity. Cell Press, 50(2), pp. 378-389.e5. doi: 10.1016/J.IMMUNI.2019.01.009.

Dyer, Douglas P, Medina-Ruiz, L., Bartolini, R., Schuette, F., Hughes, C. E., Pallas, K., Vidler, F., Macleod, M. K. L., Kelly, C. J., Lee, K. M., Hansell, C. A. H. and Graham, G. J. (2019) ‘Chemokine Receptor Redundancy and Specificity Are Context Dependent’, Immunity, 50(2), pp. 378-389.e5. doi: https://doi.org/10.1016/j.immuni.2019.01.009.

Gouon-Evans, V., Rothenberg, M. E. and Pollard, J. W. (2000) ‘Postnatal mammary gland development requires macrophages and eosinophils’, Development, 127(11), pp. 2269 LP – 2282. Available at: http://dev.biologists.org/content/127/11/2269.abstract.

Hassiotou, F., Hepworth, A. R., Metzger, P., Tat Lai, C., Trengove, N., Hartmann, P. E. and Filgueira, L. (2013) ‘Maternal and infant infections stimulate a rapid leukocyte response in breastmilk’, Clinical & translational immunology. Nature Publishing Group, 2(4), pp. e3–e3. doi: 10.1038/cti.2013.1.

Jamieson, T., Cook, D. N., Nibbs, R. J. B., Rot, A., Nixon, C., Mclean, P., Alcami, A., Lira, S. A., Wiekowski, M. and Graham, G. J. (2005) ‘The chemokine receptor D6 limits the inflammatory response in vivo’, Nature Immunology, 6(4), pp. 403–411. doi: 10.1038/ni1182.

Jäppinen, N., Félix, I., Lokka, E., Tyystjärvi, S., Pynttäri, A., Lahtela, T., Gerke, H., Elima, K., Rantakari, P. and Salmi, M. (2019) ‘Fetal-derived macrophages dominate in adult mammary glands’, Nature Communications, 10(1), p. 281. doi: 10.1038/s41467-018-08065-1.

Kwok, M. K., Leung, G. M., Lam, T. H. and Schooling, C. M. (2011) ‘Early Life Infections and Onset of Puberty: Evidence From Hong Kong’s Children of 1997 Birth Cohort’, American Journal of Epidemiology, 173(12), pp. 1440–1452. doi: 10.1093/aje/kwr028.

Lilla, J. N. and Werb, Z. (2010) ‘Mast cells contribute to the stromal microenvironment in mammary gland branching morphogenesis’, Developmental Biology, 337(1), pp. 124–133. doi: https://doi.org/10.1016/j.ydbio.2009.10.021.

MM Richert, KL Schwertfeger, JW Ryder, S. A. (2000) ‘An atlas of mouse mammary gland development.’, Journal of Mammary Gland Biology and Neoplasia, 2, pp. 227–241. Available at: https://link.springer.com/article/10.1023/A:1026499523505.

Nagarajan, D. and McArdle, S. E. B. (2018) ‘Immune Landscape of Breast Cancers’, Biomedicines, 6.

Nazari, S. S. and Mukherjee, P. (2018) ‘An overview of mammographic density and its association with breast cancer’, Breast cancer (Tokyo, Japan). 2018/04/12. Springer Japan, 25(3), pp. 259–267. doi: 10.1007/s12282-018-0857-5.

Nibbs, R. J. B. and Graham, G. J. (2013) ‘Immune regulation by atypical chemokine receptors’, Nature Reviews Immunology. Nature Publishing Group, a division of Macmillan Publishers Limited. All Rights Reserved., 13, p. 815. Available at: https://doi.org/10.1038/nri3544.

Plaks, V., Boldajipour, B., Linnemann, J. R., Nguyen, N. H., Kersten, K., Wolf, Y., Casbon, A.-J., Kong, N., van den Bijgaart, R. J. E., Sheppard, D., Melton, A. C., Krummel, M. F. and Werb, Z. (2015) ‘Adaptive Immune Regulation of Mammary Postnatal Organogenesis’, Developmental Cell. Elsevier, 34(5), pp. 493–504. doi: 10.1016/j.devcel.2015.07.015.

Pollard, J. W. and Hennighausen, L. (1994) ‘Colony stimulating factor 1 is required for mammary gland development during pregnancy’, Proceedings of the National Academy of Sciences, 91(20), pp. 9312 LP – 9316. doi: 10.1073/pnas.91.20.9312.

Sasmono, R. T., Oceandy, D., Pollard, J. W., Tong, W., Pavli, P., Wainwright, B. J., Ostrowski, M. C., Himes, S. R. and Hume, D. A. (2003) ‘A macrophage colony-stimulating factor receptor–green fluorescent protein transgene is expressed throughout the mononuclear phagocyte system of the mouse’, Blood, 101(3), pp. 1155 LP – 1163. doi: 10.1182/blood-2002-02-0569.

Tuaillon, E., Viljoen, J., Dujols, P., Cambonie, G., Rubbo, P.-A., Nagot, N., Bland, R. M., Badiou, S., Newell, M.-L. and Van de Perre, P. (2017) ‘Subclinical mastitis occurs frequently in association with dramatic changes in inflammatory/anti-inflammatory breast milk components’, Pediatric Research, 81(4), pp. 556–564. doi: 10.1038/pr.2016.220.

Wilson, G. J., Fukuoka, A., Love, S. R., Kim, J., Pingen, M., Hayes, A. J. and Graham, G. J. (2020) ‘Chemokine receptors coordinately regulate macrophage dynamics and mammary gland development’, Development, 147(12), p. dev187815. doi: 10.1242/dev.187815.

Wilson, G. J., Hewit, K. D., Pallas, K. J., Cairney, C. J., Lee, K. M., Hansell, C. A., Stein, T. and Graham, G. J. (2017) ‘Atypical chemokine receptor ACKR2 controls branching morphogenesis in the developing mammary gland’, Development (Cambridge), 144(1). doi: 10.1242/dev.139733.

Wiseman, B. S. and Werb, Z. (2002) ‘Stromal Effects on Mammary Gland Development and Breast Cancer’, Science, 296(5570), pp. 1046 LP – 1049. doi: 10.1126/science.1067431.

