## Supplemental Figures for "Diverse myeloid cells are recruited to the developing and inflamed mammary gland"

**Supplemental Information**

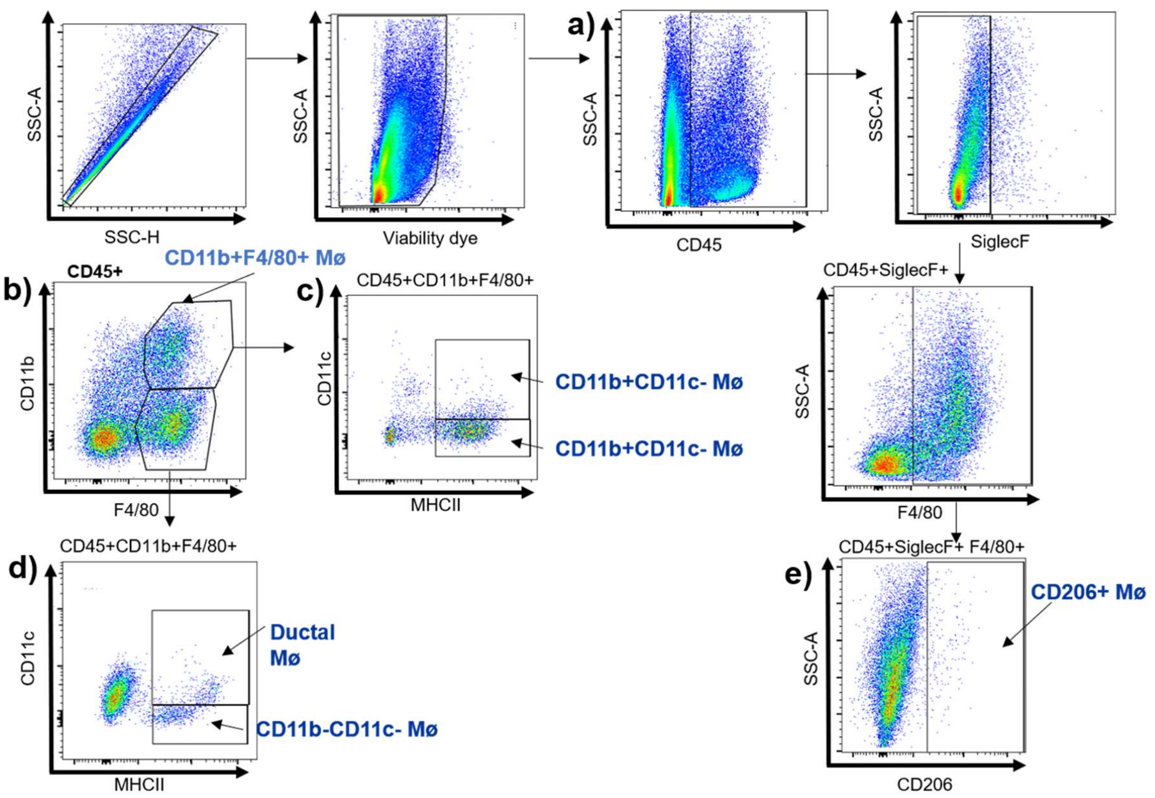


**Supplemental Figure 1: Gating strategy to define macrophage subsets in the mammary gland.** Flow cytometry was carried out to identify cell types in the mammary gland. Single cells were gated, dead cells were excluded, and **a)** CD45+ immune cells were identified. Populations were then expressed as a percentage of CD45+ cells, including; **b)** CD11b+F4/80+ macrophages, **c)** CD11b+F4/80+CD11c+ and CD11b+F4/80+CD11c+ macrophages, **d)** CD11b-F4/80+CD11c+ ductal macrophages and CD11b-F4/80+CD11c- macrophages, and **e)** SiglecF-F4/80+CD206+ macrophages.

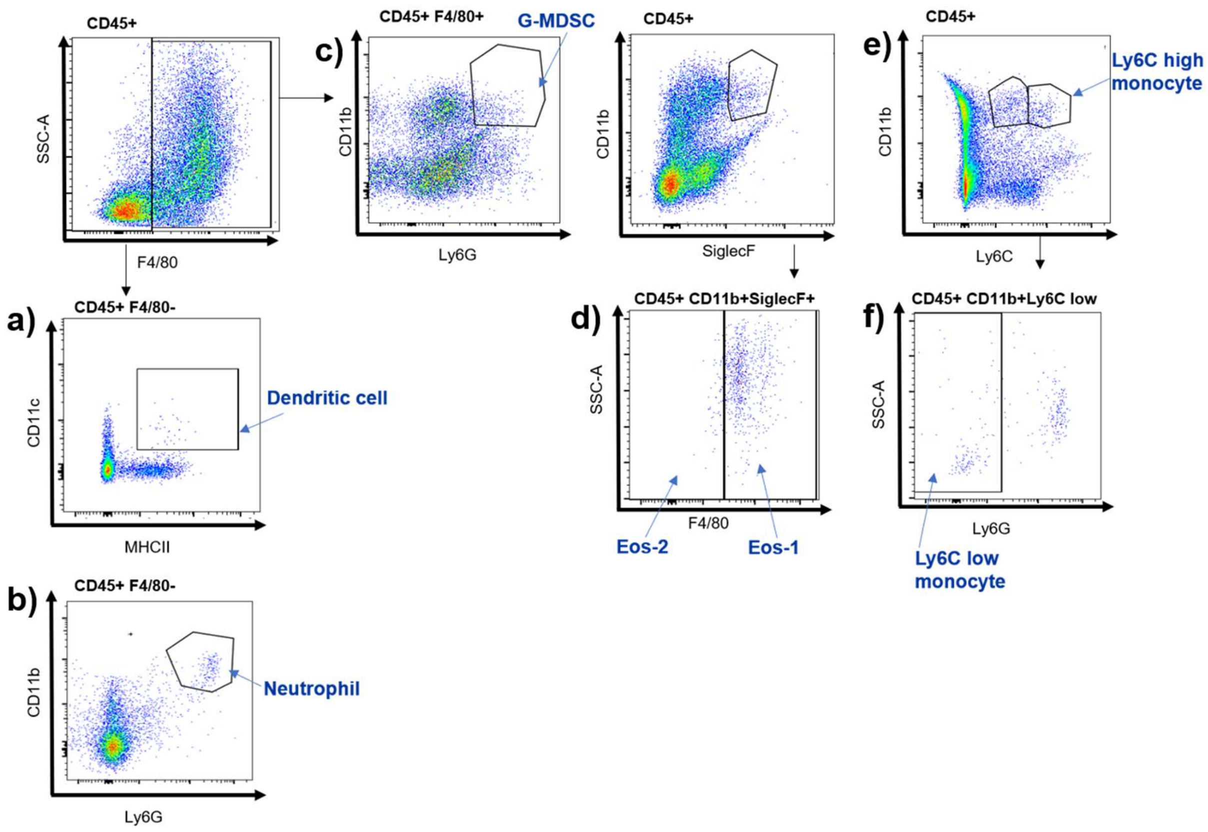


**Supplemental Figure 2: Gating strategy to define myeloid cells in the mammary gland.** Flow cytometry was carried out to identify cell types in the mammary gland. Single cells were gated, dead cells were excluded, and CD45+ immune cells were identified. Populations were then expressed as a percentage of CD45+ cells, including; **a)** F4/80-CD11c+MHCII+ dendritic cells, **b)** F4/80-CD11b+Ly6G+ neutrophils, **c)** F4/80+CD11b+Ly6G+ granulocytic myeloid derived suppressor cells, **d)** type 1 (F4/80+) and type 2 (F4/80-) eosinophils, **e)** CD11b+Ly6C high monocytes and **f)** CD11b+Ly6C low Ly6G- monocytes.

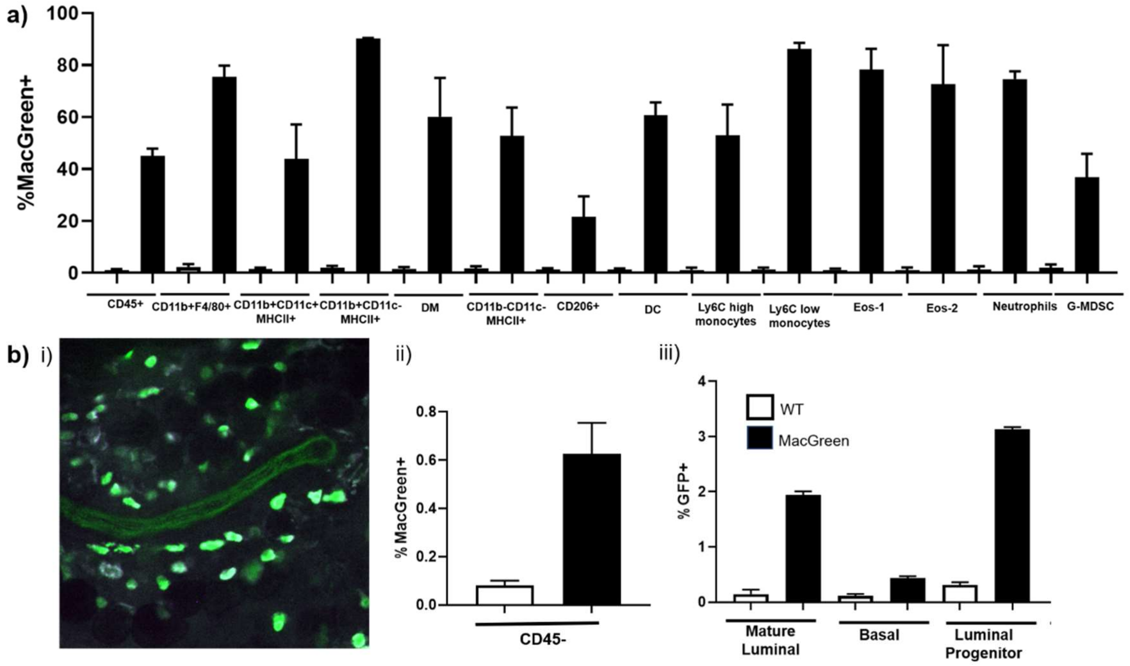


**Supplemental Figure 3: CSF1R expression in the mammary gland. a)** Flow cytometry of WT and *MacGreen* (CSF1R GFP reporter) transgenic mice to reveal the percentage of CSF1R+ cells within each cell type (n=5 per group). **b) i)** Representative confocal microscope image (25x) of a pubertal mammary gland from a *MacGreen* mouse. CSF1R+ cells are green, CD45+ cells are blue, CSF1R+ CD45+ cells are white. **ii)** Flow cytometry of WT and *MacGreen* mice to reveal the percentage of CD45- CSF1R+ cells (n=5 per group), and **iii)** within epithelial cell subsets; mature luminal (EpCAM+ CD49f-), basal (EpCAM - CD49f+) and luminal progenitors (EpCAM+ CD49f+) (n=3 per group). Error bars represent S.E.M.
